# Prospective Representations in Rat Orbitofrontal Ensembles

**DOI:** 10.1101/2020.08.27.268391

**Authors:** Jingfeng Zhou, Wenhui Zong, Chunying Jia, Matthew P.H. Gardner, Geoffrey Schoenbaum

**Author notes:** Correspondence or requests for material should be addressed to J.Z. or G.S.

## Abstract

The orbitofrontal cortex (OFC) has been proposed to encode expected outcomes, which is thought to be important for outcome-directed behavior. However, such neural encoding can also often be explained by the recall of information about the recent past. To dissociate the retrospective and prospective aspects of encoding in the OFC, we designed a non-spatial, continuous, alternating odor-sequence task that mimicked a continuous T-maze. The task consisted of two alternating sequences of four odor-guided trials (2 sequences × 4 positions). In each trial, rats were asked to make a “go” or “no-go” action based on a fixed odor-reward contingency. Odors at both the first and last positions were distinct across the two sequences, such that they resembled unique paths in the past and future, respectively; odors at positions in between were the same and thus resembled a common path. We trained classifiers using neural activity to distinguish between either sequences or positions and asked whether the neural activity patterns in the common path were more like the ones in the past or the future. We found a proximal prospective code for sequence information as well as a distal prospective code for positional information, the latter of which was closely associated with rats’ ability to predict future outcomes. This study demonstrates a prospective behaviorally-relevant predictive code in rat OFC.

## Introduction

The orbitofrontal cortex (OFC) signals expected outcomes, which is believed to be fundamentally important for outcome-directed behavior (Rudebeck & Murray, 2014; Stalnaker, Cooch, & Schoenbaum, 2015; Wallis, 2011). The vast majority of electrophysiological evidence supporting this assertion comes from behavioral settings where different cues predict reward outcomes with different sizes, identities, probabilities, and delays, etc (Kepecs, Uchida, Zariwala, & Mainen, 2008; Klein-Flugge, Barron, Brodersen, Dolan, & Behrens, 2013; Padoa-Schioppa & Assad, 2006; Roesch, Taylor, & Schoenbaum, 2006; Tremblay & Schultz, 1999). Meanwhile, the past sensory cues, rewards, spatial directions, and behavioral choices are also reported to be reflected in the OFC neural activities (Feierstein, Quirk, Uchida, Sosulski, & Mainen, 2006; Kennerley, Behrens, & Wallis, 2011; Nogueira et al., 2017; Riceberg & Shapiro, 2017; Saez, Saez, Paton, Lau, & Salzman, 2017; Young & Shapiro, 2011; Zhou, Jia, Feng, Bao, & Luo, 2015). Such findings have been taken as evidence that the OFC, together with contributions from other interconnected brain regions such as the hippocampus, might provide a neural mechanism with which animals could mentally travel through a task model in time and recall the past events and simulate future outcomes (Behrens et al., 2018; Wang, Schoenbaum, & Kahnt, 2020; Wikenheiser & Schoenbaum, 2016; R. C. Wilson, Takahashi, Schoenbaum, & Niv, 2014).

However, from studies where different past cues or episodes lead to different future outcomes, it is not clear whether the neural patterns observed are representing the future versus simply providing a record of past events. Additionally, in most behavioral settings, future events consist of rewards, whereas past events are normally intrinsically-neutral sensory cues. As a result, prioritized reward value processing would lead to a biased finding of stronger prospective coding (Wallis, 2007; Xie, Nie, & Yang, 2018).

In the present study, we resolved these confounds by recording single-unit activity in the OFC of rats performing a non-spatial, continuous odor sequence task, conceptually similar to a continuous T-maze. Combining both single-unit and neural ensemble analyses, we tested whether the OFC neural ensemble patterns during the overlapping paths in the “virtual” T-maze task resemble neural activities that occur in the past or in the future.

## Results

### Continuous, alternating odor-sequence task

The behavioral task developed in this study was designed to mimic the continuous T-maze alternation task that has been commonly used to assess spatial memory (Greene & Naranjo, 1986; Verma & Moghaddam, 1996). Rather than moving through a sequence of locations in space, subjects moved through a sequence of odors. On each trial in the odor sequence (Figure 1A), the rats were presented with the appropriate odor at a central port and then had to decide whether to respond for a sucrose reward by poking into a fluid well (“Go”) or to withhold responding on non-rewarded trials (“No-Go”). The decision to respond for reward could be made correctly based simply on odor identity, or by using information available from the sequence. The task used 6 different odors, arranged in two 4-odor sequences: S1 and S2 (Figure 1B). The 4 odors in each sequence were designated as 4 positions (P1 – P4).

**Figure 1.**
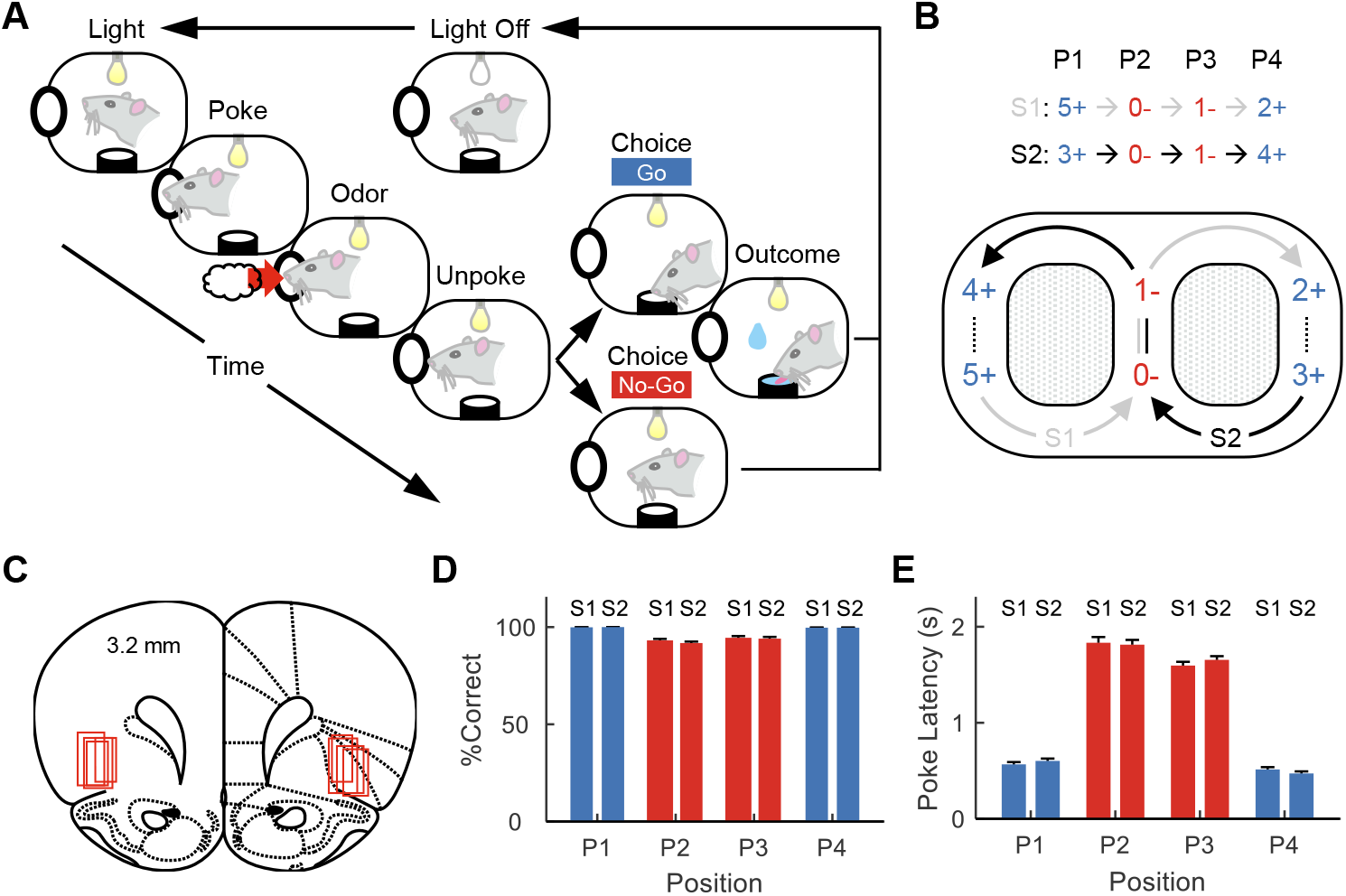
Task design, histology, and behavioral performance. (A) Single trial of the behavioral task. Rats sampled one of 6 odors from an odor port on each trial and made a “Go” choice by poking into a nearby fluid well or a “No-Go” choice by withholding their responses. (B) The 6 odors were organized into two 4-odor sequences, named S1 and S2. The four odors in each sequence represent four positions (P1 – P4). S1 and S2 alternated like a “figure eight”. (C) Reconstruction of recording sites. Red squares indicate locations of electrodes. (D) Percent correct (%correct) on each trial type during single-unit recording sessions. Blue indicates trial types with reward, while red indicates trial types without reward. Error bars are standard errors of the mean (SEMs). A two-way ANOVA (n = 64 sessions) with factors sequence (F_1,504_ = 1.03; p = 0.31; η^2^ = 0.0014) and position (F_3,504_ = 84.0; p = 4.3×10^−44^; η^2^ = 0.33) was performed. No significant interaction was observed (F_3,504_ = 0.66; p = 0.57; η^2^ = 0.0026). (E) Poke latency measures the time from light onset to odor port entry. Error bars are SEMs A two-way ANOVA (n = 64 sessions) with factors sequence (F_1,504_ = 0.10; p = 0.75; η^2^ = 4.0×10^−5^) and position (F_3,504_ = 681.0; p =8.0×10^−177^; η^2^ = 0.8) was performed. No significant interaction was observed (F_3,504_ = 0.79; p = 0.5; η^2^ = 9.3×10^−4^).

After training in this simple task, we recorded single-unit activity bilaterally from the lateral OFC (n = 1568 neurons; 4 rats). During recording, the rats performed the task with high accuracy as assessed by percent correct (%correct; Figure 1D) on each of the 8 trial types. Importantly, we also found the time rats spent to initiate a trial (i.e., poke latency; Figure 1E) was different depending on reward availability on current trials, suggesting the rats were using information available from the sequence structure to make predictions about future outcomes that influenced their responses.

As in a continuous T-maze alternation task, each position shares the same past, current, and future reward structure, thereby eliminating any bias that value might have on prospective versus retrospective encoding. Further, differences in sensory experience are structured, so that some positions (P1 and P4) differ locally, since the odors are unique at these positions in S1 and S2, while sharing the same recent past and future events (they come from and return to P2 and P3, each of which share a single odor cue in S1 and S2), whereas other positions (P2 and P3) are similar locally, since the local odors are the same, but differ in recent past and future events (they come from and go to P1 and P4, each of which have different odor cues in each sequence). This arrangement provides a unique opportunity to dissociate retrospective versus prospective neural representations. Specifically, activity distinguishing the sequences should weaken across P2 and P3 if it is retrospective, while it should grow stronger if it is prospective. Further retrospective activity might resemble neural activity patterns from the past (P1), while prospective activity might resemble activity in the future (P4). We analyzed activity recorded in OFC in well-trained rats performing this task to test these predictions.

### Distinguishing odor sequences by OFC single-units and neural ensembles

We first examined OFC neural activity at the single-unit level. We found some neurons showing differential responses to sequences S1 and S2 at all 4 positions (Figure 2A-D). A selectivity analysis indicated that the neurons showed the most selectivity at P1 and P4 around odor sampling, with fewer neurons showing selectivity at P2 and P3 (Figure 3A). Indeed, the number of selective neurons was above the chance level (5%) only around the odor for P2 and after the outcome for P3.

**Figure 2.**
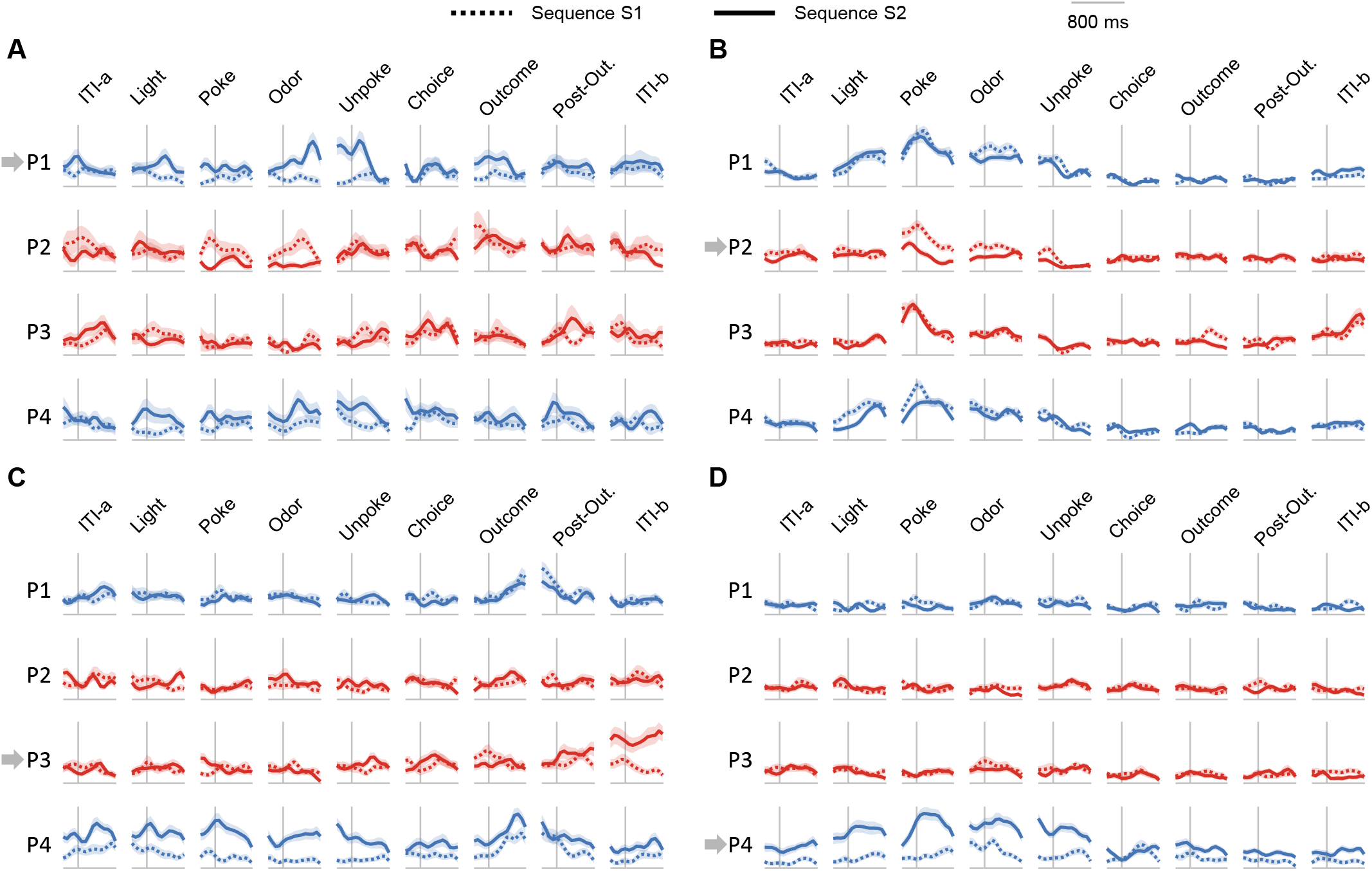
Example of single-units. (A-D) Four example neurons that exhibited differential firing to sequences S1 versus S2 at different positions (P1 – P4, indicated by grey arrows). Blue and red colors mean reward and non-reward trial types, respectively. Shaded areas indicate SEMs.

**Figure 3.**
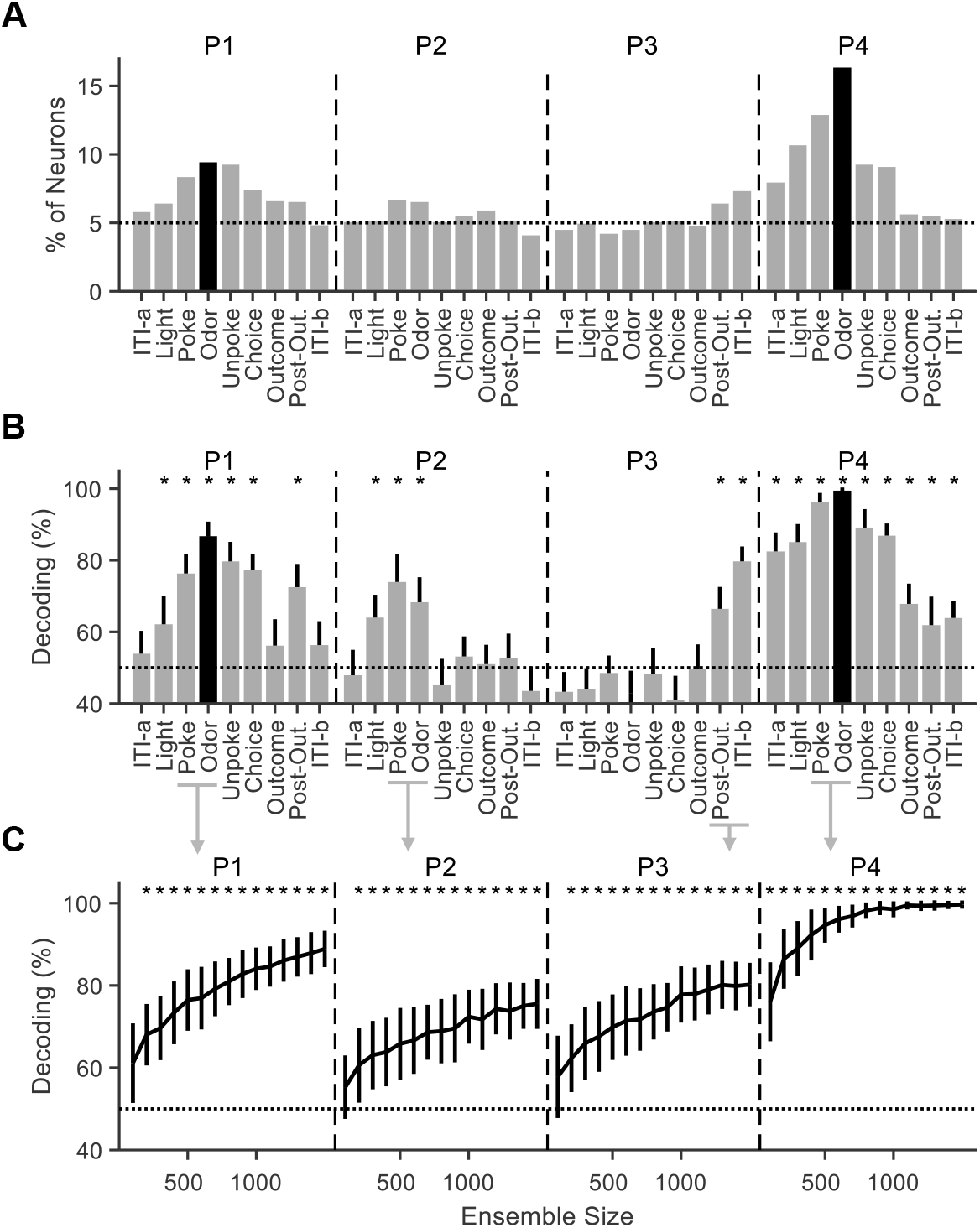
Discriminating sequences S1 versus S2 at both the single-unit and neural ensemble levels. (A) The neuronal selectivity to sequences S1 versus S2 of each neuron was calculated at different epochs [“ITI-a”, “Light”, “Poke”, “Odor”, “Unpoke”, “Choice”, “Outcome”, “postOut. (postOutcome)”, “ITI-b”] for all the four positions (p < 0.05; One-way ANOVA). The p values were not corrected; the dotted lines indicate the chance or baseline level of selectivity given this criterion. Note that only at two task epochs [“Odor” at P3 and P4; black bars] could the rats discriminate the two sequences (S1 versus S2) only based on current sensory information. At all other task epochs, shown as grey bars, an internal memory of the sequences had to be used. (B) Accuracy of decoding S1 versus S2 in each of the 9 task epochs within each position (P1 – P4). Error bars are standard deviations (SDs) and each asterisk indicates that the mean decoding accuracy exceeds a 95% confidence interval (CI) calculated using the same decoding process but with label-shuffled data. The meaning of black bars is the same as in panel A. (C) Accuracy of decoding S1 versus S2 at each position with different ensemble sizes. Task epochs used for P1, P2 and P4 are “Poke” and “Odor”, while task epochs used for P3 are “Post-Out.” and “ITI-b”. Error bars show SDs and the asterisks indicate that the mean decoding accuracy exceeds 95% CIs as results of decoding with shuffled trial labels.

Next, we examined the ability of the pattern of activity across all recorded neurons to decode sequences (S1 vs. S2) within the individual task epochs at each position. Consistent with the single-unit selectivity analysis (Figure 3A), decoding accuracy was the highest at P1 and P4, while at P2, the highest decoding happened before and after the odor delivery, and at P3, the highest decoding happened after the outcome (Figure 3B-C).

### A proximal prospective code about odor sequences

To study whether the neural patterns during the delay epochs (epochs after P1 odor time and before P4 odor time) resembled the past (S1 vs. S2; odors 5+ vs. 3+ at P1) or the future (S1 vs. S2; odors 2+ vs. 4+ at P4), we trained linear support vector machine (SVM) classifiers to distinguish the sequences during the odor period at either P1 or P4 (i.e., retrospective and prospective templates; Figure 4A-B) and then used each classifier to decode the neural activity patterns in all task epochs. This analysis revealed chance decoding at most points at the delay epochs, particularly for decoding by the classifier trained with the retrospective template, which dropped to chance immediately after odor sampling in P1 (Figure 4A). Decoding by the prospective template was also at chance for most of the delay epochs, however it increased rapidly at the inter-trial interval (ITI) and initial epochs of the P4 trials before the odor was presented (Figure 4B), suggesting the emergence of a prospective representation of the impending odors.

**Figure 4.**
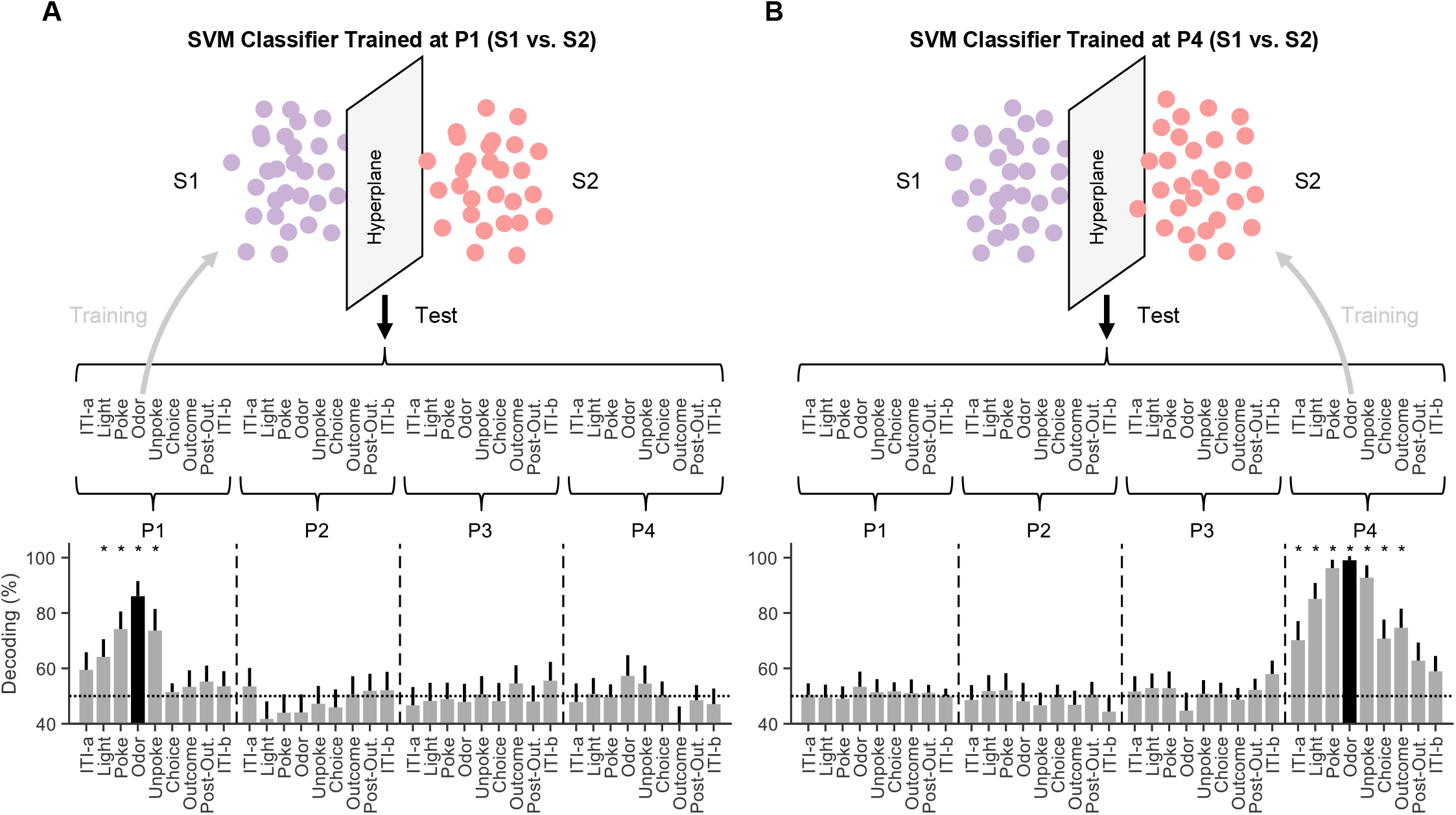
Cross-epoch decoding of sequences. (A) Each dot represents a single trial in the high-dimensional ensemble activity space (n = 30 trials for each trial type). A binary linear SVM classifier was trained to discriminate sequences S1 vs. S2 by using neural activities during odor sampling during P1 (30 trials for each trial type). The trained classifier was used to test how well S1 vs. S2 could be decoded by using the neural ensemble activities at all task epochs (9 epochs within each positions; P1 – P4). Error bars show SDs and asterisks over bar plots indicates that the mean decoding accuracy exceeds 95% CIs calculated using the same decoding process but with label-shuffled data. (B) The same as in (A) except that the SVM classifier was trained by using neural activities at P4 odor time.

### A distal prospective code to reflect future positions

A prospective code distinguishing the odor *sequences* was apparent in the run up to the odor in P4, however the prior analysis found no evidence of a stable, sustained prospective code across the entire delay epoch. This is not completely surprising because, as we previously reported, sequences in an odor sequence task tend to be generalized in the OFC if distinguishing them is not necessary for the rats to correctly perform the task (Zhou et al., 2019). This is even true when the odors differ at a given position, thus the failure of OFC to distinguish the sequences during P2 and P3 here is consistent with that prior data.

However the *positional* information was task-relevant since it was used by rats to calculate their current distance to future reward, evidenced by their different poke latencies on trial types with different probabilities of reward (Figure 1E). Thus we next asked whether we could find a prospective code for this *positional* information (i.e., a mental simulation of future epochs collapsing across the *sequence* information at each position) during the delay epochs.

To do this, we lumped S1 and S2 together at each task epoch and built a binary SVM classifier using neural activity from odor sampling during P1 and P4 (P1 vs. P4; odor time); then we used this classifier to reexamine how the neural activity patterns evolved during the delay epochs (Figure 5). Interestingly this analysis did not reveal a clear pattern of representation when averaged across rats, however when each rat was analyzed separately it revealed significant prospective activity in 3 subjects that was masked by retrospective activity in one subject (Figure 6A-D). Importantly, such a prospective code appeared early at P2 and P3 (Rat #2 and #3), and even at P1 (Rat #4), in a phasic but not tonic manner, which is dramatically different from the proximal prospective code about sequences as shown in Figure 4. Moreover, the emergence of a distal prospective ensemble code during the delay epochs was closely associated with rats’ poke latency; the stronger the prospective activity, the stronger the poke latency differed prior to rewarded versus non-rewarded trials for a subject (Figure 6E-H).

**Figure 5.**
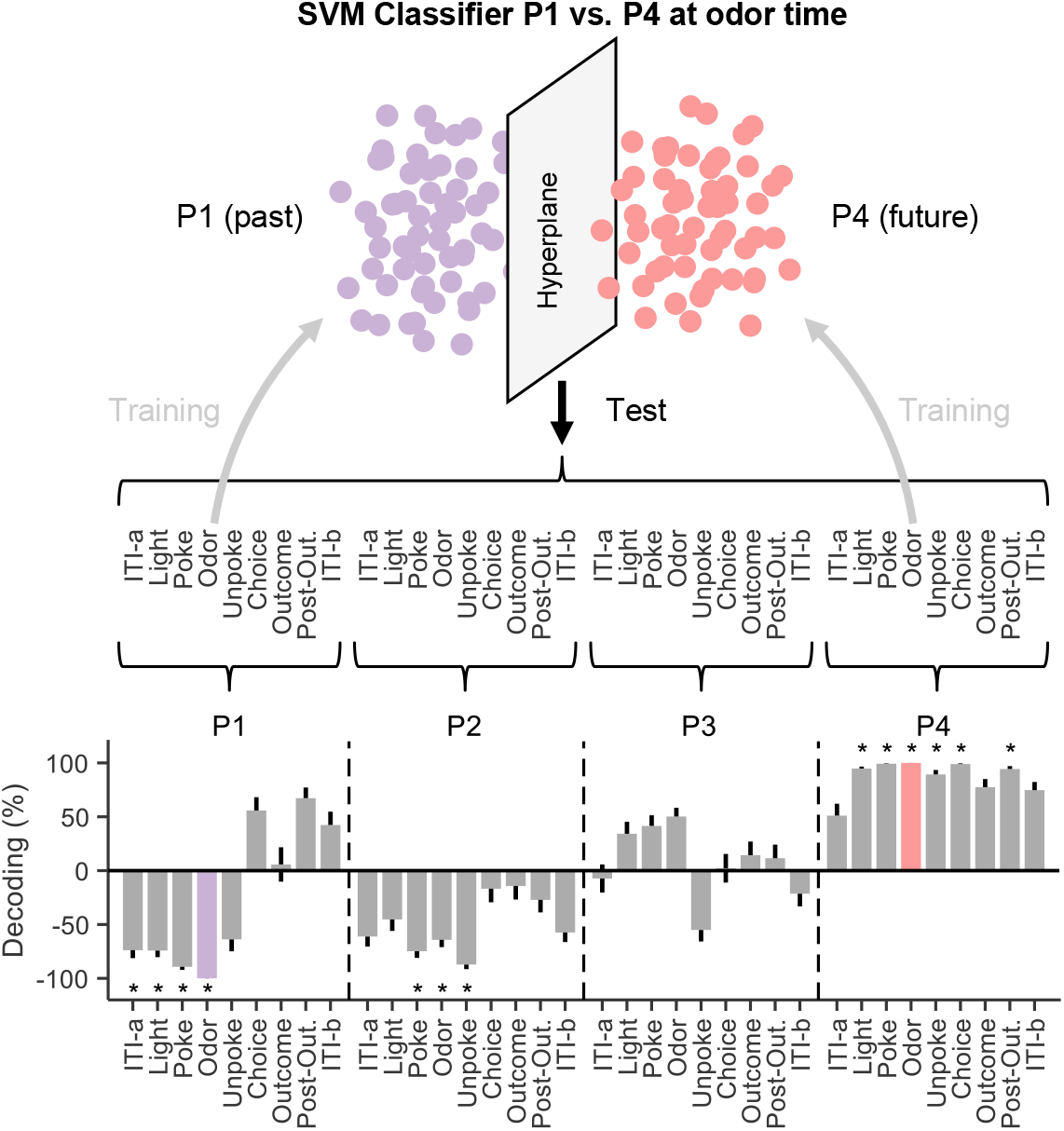
Cross-epoch decoding of positions. Trials in S1 and S2 were combined in each task epoch at 4 positions. Each dot represents a single trial in the high-dimensional ensemble activity space (n = 60 trials for each trial type). A binary linear SVM classifier was trained to discriminate positions P1 vs. P4 by using neural activities at odor sampling. The trained classifier was then used to decode neural ensemble activities at all task epochs (9 epochs within each positions; P1 – P4). Error bars show SDs and asterisks over bar plots indicates that the mean decoding accuracy exceeds 95% CIs calculated using the same decoding process but with label-shuffled data. Positive values for decoding accuracy indicate that activity at a particular position/epoch was more often classified as P4 than P1, while negative values means the opposite.

**Figure 6.**
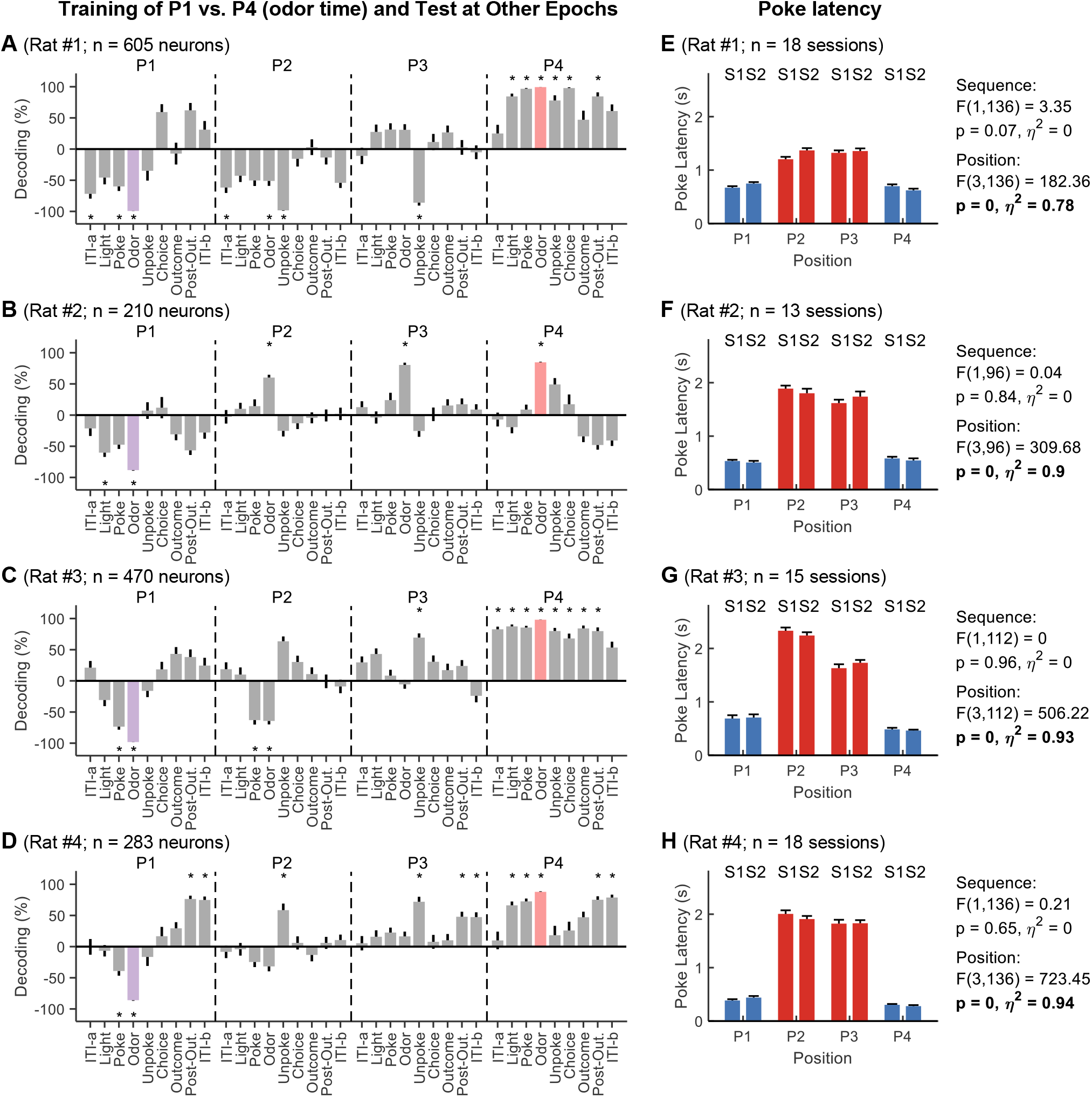
Cross-epoch decoding of positions for each rat. (A-D) Cross-epoch decoding of P1 versus P4 was performed on each rat, using the same approach and conventions as in Figure 5. (E-H) Poke latency as in Figure 1E, calculated separately for each rat. Two-way ANOVAs were performed with two factors (sequence and position). Statistical results were shown on the right side of bar plots.

To further confirm this finding, we plotted the individual sessions based on the poke latency difference between rewarded and non-rewarded trial types (Figure 7). The bimodal distribution in Figure 7A shows that the sessions with low differences (i.e. little influence of future outcome) all came from Rat #1. When these sessions were excluded, the analysis revealed a clear prospective code similar to that was seen with individual rats (Figures 7B, 6B-D).

**Figure 7.**
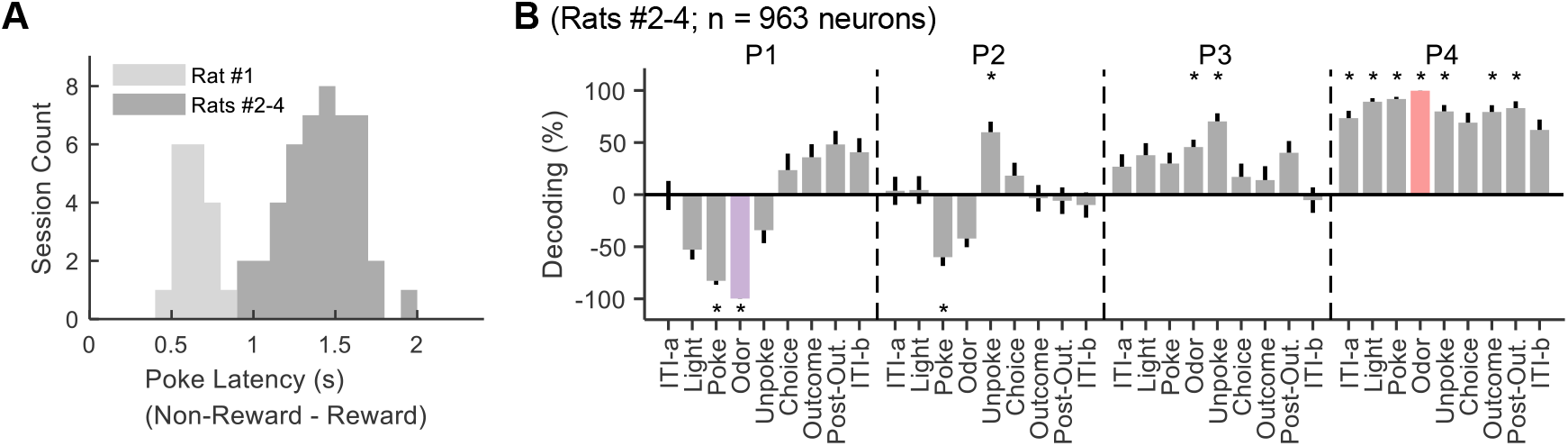
Cross-epoch decoding of positions based on performance. (A) The histogram of poke latency differences between rewarded and non-rewarded trial types. Rats #2-4 showed substantially larger differences in poke latencies than Rat #1 (B) Cross-epoch decoding of P1 versus P4 was performed on sessions from Rats #2-4, which showed large differences in their poke latencies prior to rewarded vs. non-rewarded trials, consistent with use of the sequences to predict upcoming reward. This analysis used the same approach and conventions as in Figure 5.

Together, these results suggest that both proximal and distal prospective codes exist in the OFC neural ensemble activities.

## Discussion

Through analyzing neural activities recorded in the OFCs of rats performing an odor sequence alternation task, we found two types of prospective neural ensemble codes for future events. OFC ensembles used a proximal code for immediate sequence information, while used a distal neural code for future positional information. The finding provides direct electrophysiological evidence that OFC activity anticipates the future even when confounds related to differences in past events or the prioritization of future reward information are controlled.

The stringent control on possible confounds addresses gaps in our understanding that are typically ignored. For instance, differential activity after a cue but before a reward is often taken as evidence of encoding of expected outcomes and yet it could equally well be a retrospective activity, a reflection of trace memory. Further, even if multiple cues are used to predict the same reward, the value asymmetry could lead to a bias toward apparently prospective representation even where a functional bias does not exist. The task used here, though simple to perform and perhaps a bit boring even, resolves these confounds, allowing us to discern activity that is clearly prospective.

The distal prospective ensemble code is particularly intriguing because it seems to fit well with a role in prospective memory (“remembering to remember”), a higher-order brain function found in humans, non-human primates, and rodents (Beran, Evans, Klein, & Einstein, 2012; Beran, Perdue, Bramlett, Menzel, & Evans, 2012; Evans & Beran, 2012; McDaniel & Einstein, 2007; A. G. Wilson & Crystal, 2012; A. G. Wilson, Pizzo, & Crystal, 2013). There are proposed to be three phases of prospective memory: activation or initial encoding -> inactivation when subjects are engaged in other irrelevant activities -> and reactivation when relevant information (event-based or time-based) is encountered (McDaniel & Einstein, 2007; A. G. Wilson & Crystal, 2012; A. G. Wilson et al., 2013). The pattern of prospective encoding identified here in OFC matches this evolution, mostly disappearing in the ITI periods and during the delay epochs except for brief activations when relevant sensory information was delivered around the odor period. This pattern illustrates that OFC is important for anticipating future events but also illustrates that active spiking in OFC is not sufficient for the memory to be maintained. OFC is likely supported in this by other brain regions, which may hold sustained or dynamic neural activation during the long delay time. However it is likely that maintaining such information across the relatively long delay period used here requires additional mechanisms such as short-term synaptic plasticity, either in OFC or elsewhere, which is not necessarily reflected in increased or decreased neural firing rates (Barbosa et al., 2020; Mongillo, Barak, & Tsodyks, 2008; Stokes, 2015). More investigations are needed in the future to shed light on this important question.

## ACKNOWLEDGMENTS

The authors thank the NIDA IRP histology core for technical assistance with histology. This work was supported by the Intramural Research Program at the National Institute on Drug Abuse (ZIA-DA000587). The opinions expressed in this article are the authors’ own and do not reflect the view of the NIH/DHHS. This work used the computational resources of the NIH HPC Biowulf cluster (http://hpc.nih.gov).

## AUTHOR CONTRIBUTIONS

J.Z. and G.S. designed the experiments; J.Z. collected the data; J.Z. analyzed the data with advice and technical assistance from W.Z., C.J., M.P.H.G.; and J.Z. and G.S. wrote the manuscript with input from the other authors.

## MATERIALS & CORRESPONDENCE

Correspondence or requests for material should be addressed to J.Z. or G.S.

## METHODS & MATERIALS

### Subjects

Subjects were 4 male Long-Evans rats (Charles River, 175 – 200 g, ~3-month-old) individually housed on a 12-h light/dark cycle and given ad libitum access to food in an animal facility at the AAALAC-accredited animal care facility at the National Institute on Drug Abuse Intramural Research Program (NIDA-IRP). Rats were water-deprived the day before any testing and received free access to water for 10 min in their home cages each afternoon after testing. If there was no testing the next day, the rats were given free access to water. All behavioral testing was carried out at the NIDA-IRP. Animal care and experimental procedures complied with the U.S. National Institutes of Health (NIH) guidelines and were approved by the Animal Care and Use Committee (ACUC) at the NIDA-IRP.

### Behavioral task

The behavioral training was conducted in aluminum boxes (~18” on a side) equipped with a port for odor delivery and a well for delivery of sucrose solution. Task events were controlled by a custom-written C++ program and a system of relays and solenoid valves; entries into the odor port and the fluid well were detected by infrared beam sensors. The availability of each trial was signaled by the illumination of two house-lights located on the wall above the odor port. Nosepoke into the odor port within 5 seconds after light onset initiated the trial, leading to odor delivery after a 500-ms delay. Rats were required to remain in the port for an additional 500-ms; the trial was aborted, and the lights extinguished if the rat left the odor port before this time had elapsed. After 500-ms, the rats were free to leave the port, which terminated odor delivery. After port exit, rats had 2-s to respond at the fluid well. On rewarded trials, a response led to the delivery of a sucrose solution (10% w/v; 50 μL) after a random delay ranging from 400 to 1500-ms. Upon exit from the well, non-responding during the 2-s period, or responding on non-rewarded trials, the house lights were extinguished, indicating the end of the trial and the beginning of the inter-trial interval (ITI). A 4-s ITI followed correct trials, and an 8-s ITI followed trials on which the rat made an error.

On each trial, one of 6 odors was delivered to the odor port. The 6 odors were organized into two sequences (S1 and S2) that occurred in turn repeatedly (S1→S2→S1→S2→…→S1→S2; 40 repeats of each sequence), described as below. The odor identity is indicated by a number, and reward and non-reward is indicated by the positive (+) and negative (−) symbols, respectively:

S1: 5+ 0− 1− 2+
S2: 3+ 0− 1− 4+

Rats were trained on the full set of sequences since Day 1 until they were able to perform accurately (> 75% correct) on every trial type in a session, then electrode arrays were implanted bilaterally in OFC.

### Surgical procedures

Rats were implanted with two drivable bundles of 16 electrodes (32 electrodes in total), made from nickel-chromium wires (25 μm in bare diameter; AM Systems, WA) in bilateral OFCs (AP: 3 mm, ML: 3.2 mm). Each wire bundle was housed in a 27-gauge stainless-steel tubing and cut with a pair of fine spring scissors to extend 1.5 – 2 mm beyond the end of the tubing. The tips of wires were initially placed at 4.2 mm ventral from the brain surface. After surgery, rats were given Cephalexin (15 mg/kg) orally twice a day for two weeks to prevent any infection.

### Single-unit recording

Electrophysiological signals and behavioral event timestamps were recorded with the Plexon OmniPlex System (Plexon, Dallas, TX). The initial wideband signals collected by the electrodes were amplified and digitalized at 40 kHz through a digital headstage (Digital Headstage Processor; DHP) and filtered in the control software (PlexControl) to isolate spike-band frequency (250 – 8, 000 Hz) signals. Before the start of each recording session, a common median reference (CMR) for each electrode bundle (16 electrodes in each bundle; two bundles for each rat) was used to remove online noise and artifact. A threshold for each channel was set manually for each active channel to capture unsorted spikes. Spikes were sorted later offline to remove noise and isolate single units using Offline Sorter (Plexon, Dallas, TX) with a built-in template matching algorithm. Sorted files were saved as NeuroExplorer (Nex Technologies, Colorado Springs, CO) formatted files, which were exported to MATLAB (MathWorks, Natick, MA) to extract unit and behavioral event timestamps and for further analyses. Immediately after each session, the electrodes were moved 40~80 μm ventrally in order to change the neural population being sampled.

After the recording experiments, rats were euthanized by an overdose of isoflurane and perfused with phosphate-buffered saline (PBS) followed by 4% paraformaldehyde. A small constant current was passed through each of the electrode wires to mark the final locations of electrodes. Brains were cut in 40 μm for standard histological examination.

### Quantification and statistical analyses

The number of rats and neurons were not predetermined by any statistical methods but are comparable to those reported in previous publications from our lab. All data were analyzed using MATLAB (MathWorks, Natick, MA).

### Peri-event spike dynamics

Each trial was separated into 9 epochs associated with different task events: “ITI-a”, “Light”, “Poke”, “Odor”, “Unpoke”, “Choice”, and “Outcome”, “postOutcome”, “ITI-b”. “ITI-a” marked the time point 0.7 s before the house-light on. On reward trials, the time of well-entry was labeled as “Choice”. “Outcome” was at the time of reward delivery. On non-reward trials, the end of the 2-second window for responding was labeled as “Choice” and a time point 0.7 s after the “Choice” as “Outcome”. On both reward and non-reward trials, 0.7 s after the outcome was labeled as “postOutcome”, and 0.7 s after that was labeled as “ITI-b”. Behavioral performance was quantified by the percent of trials on which the rats responded correctly and the latency with which they initiated a trial after light onset. The spike train for each isolated single unit was aligned to the onset of each task event for the calculation of a peri-event time histogram (PETH). Pre-event time was set to be 200 ms, and post-event time 600 ms. Spike number was counted with a bin = 100 ms. A Gaussian kernel (σ = 50 ms) was used to smooth the PETH on each trial.

For further analyses, only 30 correct trials were randomly selected from each trial type (30 trials × 8 trial types = 240 trials in total); and the post-event firing rates (100 – 600 ms) were averaged to obtain a single measurement of neural activity for each neurons on each trial at each task epoch.

### Classification analyses

The neural data at each task epoch was organized as a 2-dimensional matrix (trials × neurons) in a way that each row represents one trial and each column represents the firing rates of one neuron in all the trials. In other words, each trial is a vector in which each dimension is the firing rate of one neuron. Neurons recorded from different sessions were concatenated with alignment to the trials to form pseudo-ensembles. We shuffled trial orders within each trial type to generate a different pseudo-ensemble as well as to remove the temporal correlation between neurons. The trial-order shuffling was repeated for 10, 000 times such that 10, 000 pseudo-ensembles were generated.

We used the linear support vector machine (SVM) for classification analyses (Chang & Lin, 2011). The classification accuracy was assessed by a leave-one-out cross-validation procedure. Specifically, one trial from each trial type was left out for future testing, and all the other trials were used to train the classifier. For each pseudo-ensemble, the leave-one-out cross-validation was repeated 200 times to estimate a mean decoding accuracy. The decoding analyses were carried out on the 10, 000 pseudo-ensembles to obtain an overall mean decoding accuracy. The statistical significance of the overall mean decoding accuracy was determined by the 95% confidence interval estimated by running the same decoding process with label-shuffled pseudo-ensembles. For cross-epoch classification analyses, we followed the same procedure but with training and test sets from different task epochs.

